# High throughput imaging of nanoscale extracellular vesicles by scanning electron microscopy for accurate size-based profiling and morphological analysis

**DOI:** 10.1101/2021.01.20.427457

**Authors:** Sara Cavallaro, Petra Hååg, Kristina Viktorsson, Anatol Krozer, Kristina Fogel, Rolf Lewensohn, Jan Linnros, Apurba Dev

## Abstract

Nanoscale extracellular vesicle (EVs) have been found to play a key role in intercellular communication, offering opportunities for both diagnostics and therapeutics. However, lying below the diffraction limit and also being highly heterogeneous in their size, morphology and abundance, these vesicles pose significant challenges for their physical characterization. Here, we present a direct visual approach for their accurate morphological and size-based profiling by using scanning electron microscopy (SEM). To achieve that, we methodically examined various process steps and developed a protocol to improve the throughput, conformity and image quality while preserving the shape of EVs. The investigation was performed with small EVs (sEVs) isolated from a non-small cell lung cancer (NSCLC) cell line H1975 as well as from a human serum, and the results were compared with those obtained from nanoparticle tracking analysis (NTA). While the comparison of the sEV size distributions showed good agreement between the two methods for large sEVs (diameter >70 nm), the microscopy based approach showed a better capacity for analyses on smaller vesicles, with higher sEV counts compared to NTA. In addition, we demonstrated the possibility of identifying non-EV particles based on size and morphological features. The study also showed process steps that can generate artifacts bearing resemblance with sEVs. The results therefore present a simple way to use a widely available microscopy tool for accurate and high throughput physical characterization of EVs.

## INTRODUCTION

During the past decade, research efforts have significantly enhanced our understanding of extracellular vesicles (EVs), their biological relevance and pathological significance, particularly in cancer.^1–4^ To support this rapidly expanding field, a number of analytical tools such as Nanoparticle Tracking Analysis (NTA), tunable resistive pulse sensing (TRPS), dynamic light scattering (DLS) as well as various microscopy methods including electron microscopy (EM, e.g. transmission EM, scanning EM, cryo-EM) or atomic force microscopy (AFM) has been proposed for the physical characterization of EVs.^5–7^ These methods have played a major role in the overall development of the field, as the physical characterization is a primary means to optimize various EV isolation and enrichment protocols, and benchmark them in terms of purity, concentration and size distribution of the isolated EVs. These characterization methods, however, have very large differences in their accuracy, measurable size range, throughput, analytical time and cost.^8^ While NTA, considered to be a gold standard in EV characterization, offers the benefit of high throughput and operational simplicity, the method suffers from a lower resolution and has less capacity to interrogate EVs of smaller size, i.e. below 60-70 nm.^5,8^ Besides, NTA provides only hydrodynamic radius and, thus, cannot distinguish between a spherical object and a particle of random shape.^8,9^ Moreover, the recent understanding concerning the heterogeneity in EV populations has introduced a new analytical challenge as well as a renewed interest in the high-resolution physical characterization of EVs.^10,11^ Various recent reports not only suggest that populations of EVs exist in the size range of ~50-60 nm or below,^10,12^ but also that EVs are very heterogeneous in their size and shape even when following the same route of biogenesis.^13^ In addition, recent investigations also indicate that the stiffness of EVs is influenced during certain physio-pathological conditions.^14^ This suggests that the shape/deformation/morphology of EVs are likely to become important for further understanding of their biophysical properties and functional heterogeneity. Given the inherent limitations of analysis by scattering-based approaches,^5,8^ these conditions are not likely to be fully met by the available technologies like NTA. This necessitates the development of new methodologies, which are also widely available and inexpensive. Microscopy-based approaches such as transmission EM (TEM), scanning EM (SEM) and AFM offer very high resolution, but suffer from low throughput, and low/or operator dependent quality and reproducibility.^8,15^. These drawbacks have made many of the images obtained by these techniques very heterogeneous^16–19^ or insufficient for EV size profiling.^20^ Cryo-EM, in this respect, certainly offers a major advantage, but the technique is neither widely available nor cost effective for frequent sample analysis.

In this report, we demonstrate the prospect of using scanning electron microscopy for high resolution size-based profiling of EVs. In particular, we present a comparison of different pre-imaging steps and substrate functionalization protocols for improvement in throughput and image quality. The study was performed on both cell-line derived sEVs, isolated from a non-small cell lung cancer (NSCLC) cell line H1975, and sEVs isolated from human serum. We tested different preparation protocols for SEM with NSCLC cell derived sEVs and analyzed their influence on the image quality and throughput. We then applied the optimal parameters to profile serum sEVs isolated via two different methods namely size exclusion chromatography (SEC) and tangential flow filtration (TFF). The comparison of sEV size distributions obtained by SEM and NTA revealed that while the distribution of larger sEVs (diameter > 70 nm) closely followed each other, they significantly deviated for smaller sEV sizes. The SEM based approach clearly identified a larger proportion of EV like particles in smaller size range (< 70 nm). In addition, we also identified the presence of particles of random shapes within the reported sEV size range from the analyzed samples, indicating the importance of morphological analysis for accurate EV identification and size profiling.

## MATERIALS AND METHODS

### Reagents

High purity deionized water (DIW) with a resistivity of 18 MΩ·cm was used throughout all the experiments. Phosphate-buffered saline (PBS, P4417) in tablets and a Grade I Glutaraldehyde solution (G5882) specifically purified for EM were purchased from Sigma-Aldrich. If not stated otherwise, all the other chemicals were purchased from Sigma-Aldrich and filtered using a 0.45 *μ*m filter before use.

### EV purification and isolation

The EVs investigated in this study were collected from two different sources and isolated in order to obtain small EVs (sEVs, 30-300 nm). The sEVs used to develop the SEM protocol were isolated from the cell culture media of NSCLC cell line H1975 (ATCC, LGC Standards, Germany) and isolated using IZON size exclusion chromatography columns (SEC; IZON, Oxford, UK). The H1975 cells were grown in RPMI-1640 media supplemented with exosome depleted media (Thermo Fisher Scientific). The media was collected after 48 h, was centrifuged at 200 g for 5 mins followed by 2000 g for 10 min and concentrated from 50 ml to about 500 *μ*l using Amicon® Ultra-15 Centrifugal Filter concentrators with a 3k cutoff (Merck Millipore, Solna, Sweden). The IZON EV original columns were rinsed with 15 ml PBS (0.22 *μ*m filtered) and the samples were added followed by gradual addition of PBS. Fractions of 500 *μ*l were collected and the main sEV containing fractions without protein contamination (6-10 according to the company) were pooled and concentrated using Amicon® Ultra-4 Centrifugal Filter concentrators.

The sEVs used to validate the platform were collected from a commercial human serum purchased from Merck Millipore (same lot in all preparations) and isolated using two different methods, namely SEC and tangential flow filtration (TFF). According to the manufacturer, the serum samples were filtered through a conventional 220 nm filter prior to freezing and subsequent shipment. Serum samples arrived frozen. They were thawed and used within 24 hours after thawing. For the sEVs isolated via SEC, the same protocol as the NSCLC sEVs presented above was followed. However, the serum sample was concentrated from 4 ml to around 700 *μ*l using Amicon® Ultra-4 Centrifugal Filter concentrators with a 3k cutoff. For the sEVs isolated via TFF, MicroKros filters from Spectrum Labs (now Replingen) were used in pore sizes 200 nm and ~20 nm (500 kD). Prior to use, all filters were rinsed with MilliQ water to remove glycerol, which is used by the manufacturer to preserve pore sizes (according to manufacturer description). Thereafter, filters and all tubing were blocked by a 10 mg/mL solution of bovine albumin in PBS, which was allowed to permeate through the pores for at least 3 hours, and then flushed with PBS. Serum sample of 20 mL was circulated through the MicroKros filter system for 4 h and thereafter a retentate of 4 mL was obtained, containing particles larger than about 20-30 nm (500kD) and smaller than 200 nm.

### EV characterization

The sEVs analyzed in this study were characterized by NTA for concentration and size estimation and by Western Blot (WB) for their protein characterization (CD9 and CD63). The NTA measurements were performed on a NS300 instrument (NanoSight, Malvern Panalytical, Malvern, United Kingdom). The serum sEVs were diluted 1:200 in PBS and analyzed with the following settings: syringe pump speed 100, detection threshold 8, camera level 13 and analysis time 5×60 s. The cell line sEVs were diluted 1:100 in PBS and analyzed as follows: syringe pump speed 100, detection threshold 4, camera level 14 and analysis time 3×60 s. sEVs isolated from NSCLC cells as well as from serum by SEC and TFF were analyzed for CD9 and CD63 expressions using WB. The EVs were lysed using 5×RIPA buffer, sample buffer was added, and the lysates ran on a Bis Tris gel, 4-12% with MES buffer (Thermo Fischer Scientific). The proteins were transferred to nitrocellulose membranes, blocked with 1:1 Odyssey blocking buffer: TBST, and incubated with respective primary antibody: anti-CD9 (#13403) and anti-calnexin (#2433) from Cell Signaling Technology and anti-CD63 (MAB5048) from R&D Systems (Abingdon, UK). Signals from the secondary antibody goat anti-rabbit IRDye® 800CW (LI-COR) were visualized using the Odyssey® Sa Infrared Imaging System (LI-COR).

### Functionalization protocols

For all the protocols, Si substrates (1cm × 1cm) with a thermally grown SiO_2_ layer were used. The substrates were cleaned with acetone, isopropanol and DIW in sequence. Three different functionalization protocols, as described in the following section, were investigated.

#### I. Covalent protocol

The SiO_2_ substrates were functionalized using our previously reported functionalization protocol up to the glutaraldehyde (GA) step.^21^ Briefly, the SiO_2_ wafers were first cleaned in a 5:1:1 solution of DIW, H_2_O_2_ and NH_4_OH (88°C, 10 min) and activated with (3-aminopropyl)triethoxysilane (APTES, 5% v/v in 95% ethanol, 10 min) and GA (1% v/v in 1× PBS, 1 h). Thereafter, EVs were covalently immobilized on top of GA for 1 h, using the GA-amine interaction. Following EV capture, the remaining GA active sites were deactivated with Tris-Ethanolamine (Tris-ETHA, 0.1 M Tris buffer and 50 mM ethanolamine, pH 9.0, 30 min) and casein (0.05% w/v in 1× PBS, 60 min). Finally, the functionalized substrates were washed with 1×PBS prior to the subsequent pre-imaging steps.

#### II. Non-covalent protocol

The SiO_2_ substrates were prepared following our previously reported functionalization protocol.^21^ Briefly, after the GA step, CD9 antibody (50 *μ*g/mL solution in 1xPBS; ab195422 from Abcam) was immobilized on top of it for 2 h. The antibody targeted the CD9 surface protein which is known to be expressed in the analyzed NSCLC cell sEVs.^21,22^ Thereafter, the remaining GA active sites were deactivated using Tris-ETHA and casein. After 1xPBS washing, the sEVs were incubated on top of the antibody-coated substrate for 1h. Following sEV binding, the substrates were washed with 1xPBS to remove the unbound vesicles.

#### III. Control protocol

The SiO_2_ substrates were prepared following all the steps of the covalent protocol except for the sEV incubation. Briefly, following GA, Tris-ETHA and casein were used to deactivate it. Finally, the functionalized substrates were washed with 1×PBS.

For the characterization of cell-line derived sEVs, the vesicles were immobilized/incubated at a concentration of 3×10^9^ particles/mL. For the characterization of serum sEVs, the vesicles were immobilized at a concentration of 1×10^10^ particles/mL. Unless stated otherwise, all the sEVs were fixed in a solution of GA/Paraformaldehyde (PFA) (0.1% GA and 2% PFA in 1×PBS) for 30 min.

### Pre-imaging steps

The pre-imaging steps consisted of sample drying, either in air or by using critical point drying (CPD), and sputtering. The influence of these pre-imaging steps was investigated in different combinations. In the case of air drying, the substrates with the immobilized sEVs were thoroughly washed with water after PBS, in order to minimize crystal formation, and left to dry in air in a fume hood. In the case of CPD, the substrates were first dehydrated with a series of ethanol washing cycles (20%, 40%, 60%, 80%, 100%, 10 min each) and then inserted into the CPD for liquid exchange (CO_2_) and drying. Depending on the availability, a E3100 CPD system from Quorum Technologies or an EM CPD300 CPD from Leica were used, keeping constant the drying parameters.

For the substrates that underwent sputtering prior to SEM imaging, the process was performed in a Polaron SC7640 Au/Pd sputter (Uppsala University). This metal layer was sputtered on top of the samples to increase the lateral conductivity for the SEM electron beam. The sputtering parameters were set in order to deposit a ~10 nm layer of Au/Pd on top of the substrates.

### SEM imaging and analysis

SEM imaging was performed in an Ultra 55 SEM microscope from Zeiss, using the Inlens detector, a working distance between 2 mm and 3 mm, and a 20 *μ*m aperture. Pixel Averaging was set as a noise reduction method. Moreover, the corners of the substrates were connected to the metallic sample holder using a slice of conductive Cu tape, in order to reduce charging effects and improve the image resolution. The SEM images were acquired at random locations on the substrates and the collected scans were examined via visual inspection for morphological analysis. For size distribution analysis, Fiji software was used.

## RESULTS

For the evaluation of various preparation, functionalization and isolation protocols, we acquired the SEM images by scanning different substrate areas and by analyzing the number, distribution and morphology of the immobilized sEVs. As presented below, we divided the analyzed parameters into two different groups, namely functionalization-related and pre-imaging parameters. The functionalization-related group includes all the steps necessary for the NSCLC sEV conjugation to the substrates, while the pre-imaging group describes the steps performed after the substrate functionalization and prior to SEM imaging. The characterization results of NSCLC sEVs isolated by SEC and the serum sEVs isolated by SEC and TFF are presented in the supplementary information (Figure S1).

### Functionalization-related parameters

To demonstrate the possibility of using SEM for size-based EV profiling and morphological analysis, it is important to identify and reduce artifacts that might come from non-EV particles, surface textures etc., ultimately resulting in erroneous measurements. Therefore, as a first step, we examined the effects of different functionalization parameters on control substrates, which were prepared by following the control protocol as indicated above. Figure 1A shows a representative image of a control substrate functionalized by using as purchased chemicals without any filtration and a standard GA solution. As visible, the substrates showed many particles in the sEV size range (50-200 nm) whose number significantly increased after the GA coating (data not shown). Figure 1B shows instead a representative image of a control substrate prepared using the control protocol up to the GA (standard solution) step (no Tris-ETHA, no casein). As presented, in both cases the controls showed particles in the EV size range that were clearly not vesicles. These artifacts would also appear in substrates with sEVs, therefore leading to erroneous counts and size distribution measurements. The effect was more pronounced for control substrates where the GA was not deactivated by Tris-ETHA and casein (Figure 1B). In this case, SEM detected many circular particles that could be mistaken as vesicles but were instead created by the GA reaction with some of the chemical compounds used (Figure 1B). On the other hand, Figure 1C shows a representative control substrate prepared following the optimized control protocol. In this case, we used a Grade I GA solution specifically purified for EM studies. Furthermore, we filtered all the chemicals prior to deposition, and we deactivated the GA active sites with tris-ETHA and casein.

**Figure 1.**
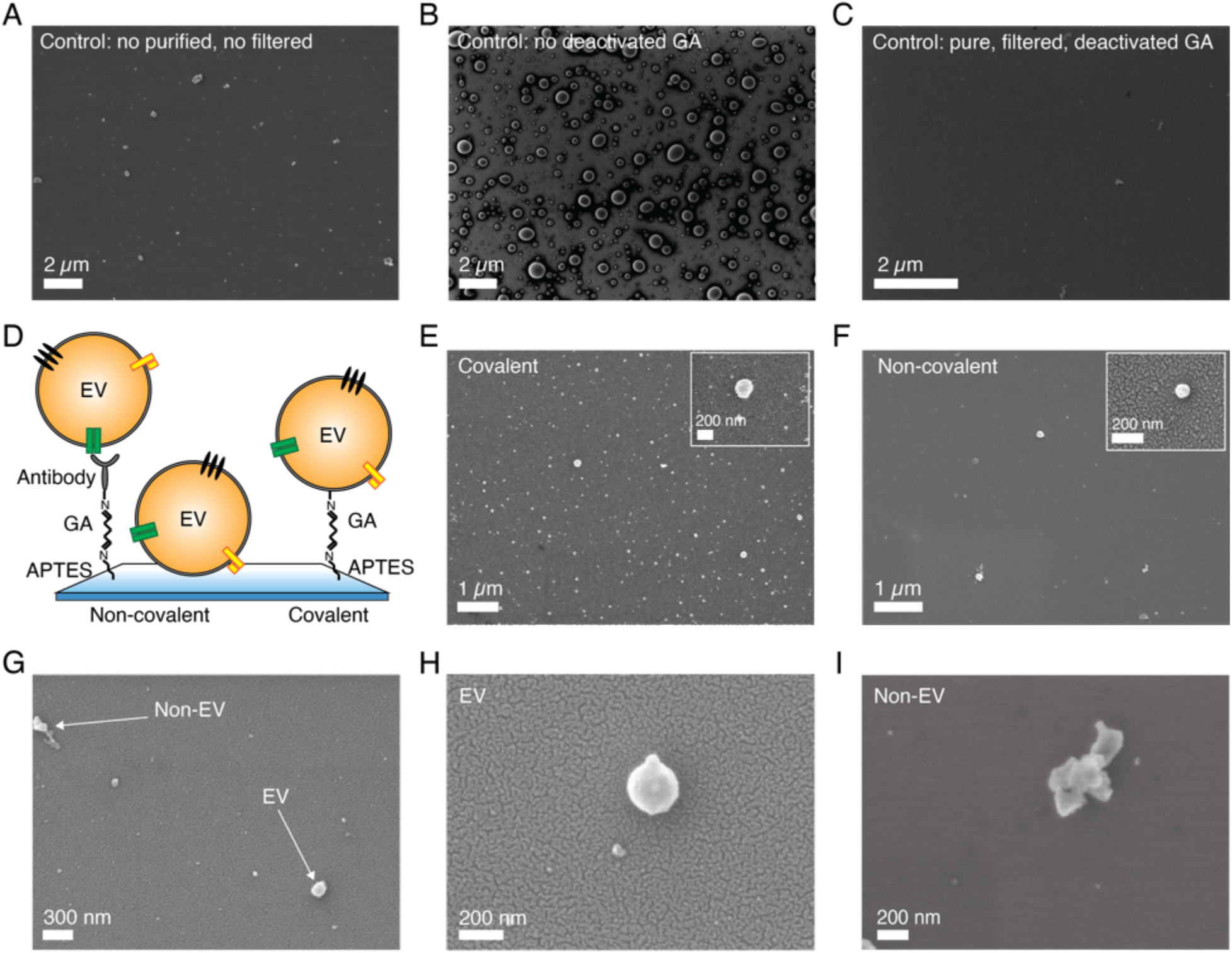
Effects of the functionalization-related parameters on the SEM imaging results for the NSCLC sEVs. (A) Control substrate prepared following the control protocol with as purchased, not filtered functionalization chemicals and a standard GA solution. Accelerating Voltage (AV)= 2 kV. (B) Control substrate prepared following the control protocol up to the GA step. No filtered chemicals deposited, and standard GA not deactivated by Tris-ETHA and casein used. AV=2 kV. (C) Control substrate prepared following the control protocol using filtered chemicals and Grade I and deactivated GA solution. AV=2 kV. No sputtering used for all control substrates in A, B, and C. (D) Schematic of sEV coupling to a SiO_2_ wafer using non-covalent and covalent captures. Non-covalent coupling performed via antibody capture (non-covalent protocol) or vesicle adsorption, while covalent capture performed via GA-amine interaction (covalent protocol). (E) Representative image of a substrate with sEVs functionalized following the protocol for covalent capture. Zoomed in image of a few vesicles detected by this strategy. AV=3 kV. (F) Representative image of a substrate with sEVs functionalized following the protocol for non-covalent capture, using anti-CD9 antibody. Zoomed in image of a vesicle detected by this strategy. AV=3 kV. (G) Representative images showing the difference between particles of spherical shape like EVs and particles of random shapes. AV=3 kV. (H) Zoomed in image of a vesicle-like particle. AV=3 kV. (I) Zoomed in image of a particle of random shape. AV=3 kV. An Au/Pd layer (~10 nm thickness) was sputtered on top of the substrates in E, F, G and H. All the samples in this figure were dried using CPD.

As presented in Figure 1C, the substrates showed the expected results, appearing very clean and with very few particles/objects on top and in the sEV size range. Figure S2 shows additional representative images of the different control substrates investigated. We emphasize that all the figures presented here depict the general features of the substrates and were verified by randomly analyzing different substrate locations (>10 spots per substrates) and also multiple substrates prepared by identical protocols. After having identified and optimized the parameters of the functionalization chemicals that lead to reliable controls, we compared the effects of covalent and non-covalent EV capture. Figure 1D schematizes these two strategies. In the case of non-covalent capture, we conjugated the vesicles to the substrate by either antibody coupling or adsorption (Figure 1D, left and center), whereas for covalent capture, the interaction between GA and amines was used (Figure 1D, right). As presented in Figures 1E, 1F and S3, the SEM results revealed that the covalent EV strategy retained a larger number of vesicles on the substrates (~8 particles/*μ*m^2^) as compared to the non-covalent one (<1 particle/*μ*m^2^). In this latter case, most of the vesicles were lost during the steps following functionalization due to the low capture strength, resulting in sparsely populated substrates in the SEM images (Figure 1F). Overall, the images showed reasonably good resolution and quality, suggesting that the analyzed sEVs had rather spherical shapes. Moreover, they confirmed the capacity of SEM to distinguish between spherical/oval objects such as vesicles and other particles of random shapes (Figures 1G, 1H, 1I). The latter could be excluded during image analysis.

### Pre-imaging parameters

As known, SEM operation requires the application of high vacuum into the microscope chamber and therefore the EVs need to be appropriately dehydrated prior to imaging, without affecting their morphology. This is a critical step for biological species, as they contain and/or are surrounded by liquid in their physiological environment. In addition, fixation might also be needed in order to preserve their shapes.^23^ Moreover, the non-conductive nature of bio-samples requires the use of a conductive layer on top of them in order to reduce charging effects arising during the electron beam-sample interaction.^24^ Therefore, following functionalization optimization, we analyzed the effects of these different pre-imaging procedures on the final SEM imaging. Figures 2 and S4 present the results of the analysis. In particular, we first analyzed and compared air drying (AD) with critical point drying (CPD), as these are the two commonly used and reported techniques in the EV field.^16–19^

**Figure 2.**
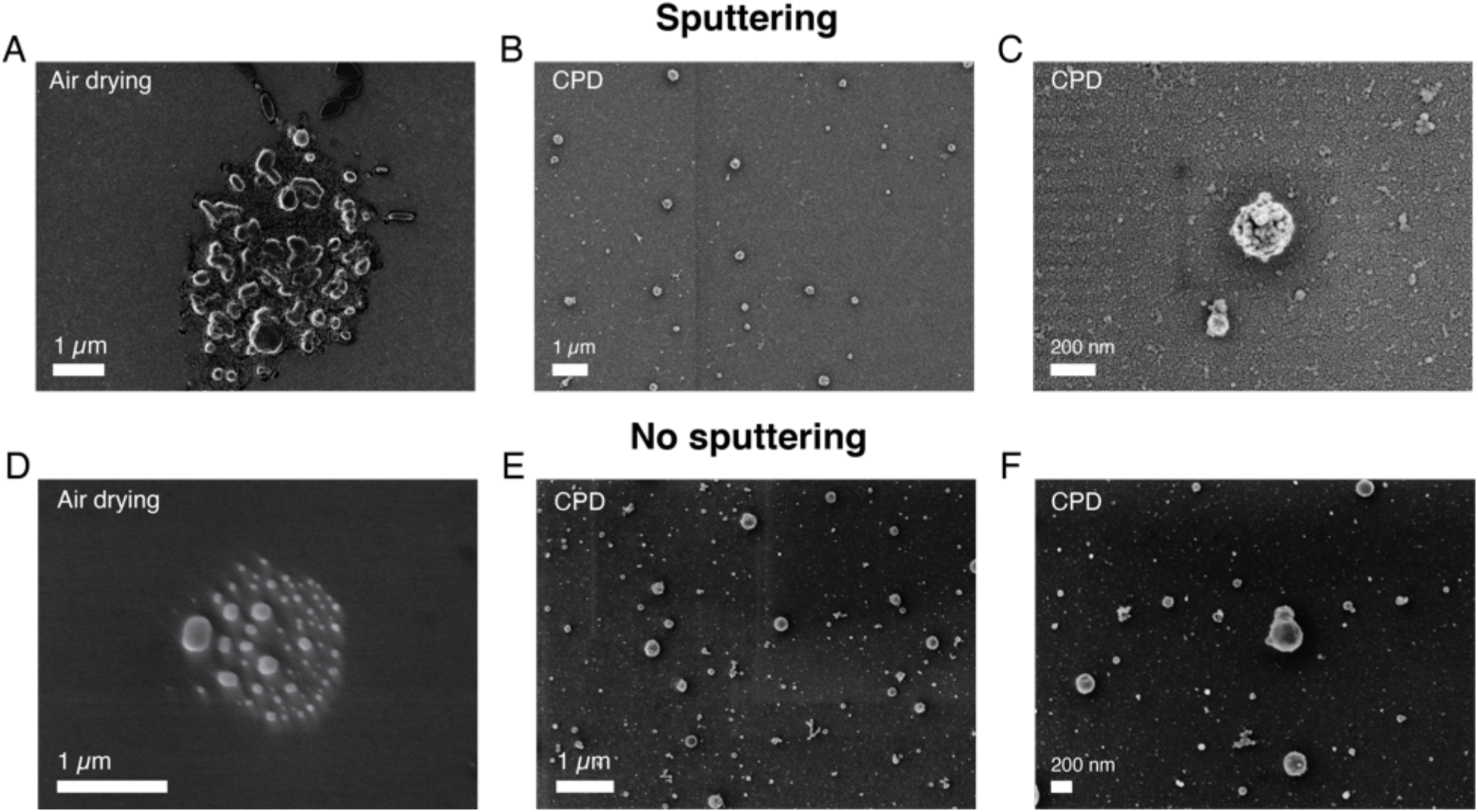
Effects of the pre-imaging parameters on the SEM imaging results for the NSCLC sEVs. (A) Representative SEM image of a substrate where the sEVs were fixed in a solution of GA/PFA and then dried in air. (B) Representative SEM image of a substrate where the sEVs were fixed in a solution of GA/PFA and then dried using CPD. (C) Zoomed in image of a few vesicles dried using CPD. An Au/Pd layer (~10 nm thickness) was sputtered on top of the substrates in A, B and C. (D) Representative SEM images of a substrate where the sEVs were fixed in a solution of GA/PFA, were dried in air but were imaged without the Au/Pd sputtered layer. (E) Representative SEM images of a substrate where the sEVs were dried using CPD but were imaged without the Au/Pd sputtered layer. (F) Zoomed in image of a few vesicles dried using CPD but imaged without the Au/Pd sputtered layer. AV=1kV for all images for better resolution of the vesicle surfaces.

Figure 2A shows a representative SEM image of sEVs that were dried in air, while Figures 2B and 2C show representative SEM images of sEVs that were dehydrated using CPD. More images comparing the two drying techniques are presented in Figure S4. As shown, in the case of CPD the vesicles appeared to be uniformly distributed on the substrate and retained their spherical shape (Figures 2B and 2C). On the contrary, the vesicles that were dried in air created a coffee stain effect,^25^ with distinct islands of clustered particles (Figures 2A and S4). Moreover, the images also suggested the presence of substrate areas (Figure S4) containing many particles with random size and shapes, and that the morphology of these vesicles was altered by the drying process. Unlike CPD processed substrates, the vesicles that were dried in air appeared to be more elongated (Figures 2A and S4). Although advantageous, CPD suffered from low reproducibility and in some cases did not show the expected results, damaging the substrates. This was possibly due to flaws during the liquid exchange and/or drying process itself. In order to reduce image blurring caused by the charging effect, we acquired the SEM images presented in Figures 2A, 2B and 2C with a thin (~10 nm) sputtered conductive layer of Au/Pd. However, this step may induce errors in size estimation, particularly for smaller vesicles. This problem can be avoided by the use of a more conductive substrate. Figure 2C shows representative SEM images obtained from sEVs that were covalently coupled to a heavily doped Si substrate. As presented, by using conductive substrates, we could still acquire good images of the vesicles, however, the resolution was slightly lower (Figure 2F) as compared to the cases where a thin metal layer was used (Figure 2C). Furthermore, while the sEVs in Figures 2B and 2C were fixed in a solution of GA/PFA, the sEVs in Figures 2E and 2F were not fixed prior to CPD. Similarly, the sEVs in Figures 2A and 2D were fixed prior to air drying, while the sEVs in Figure S5 were not. Nevertheless, in all the cases the vesicles without fixation showed similar shapes as the fixed sEVs treated with the corresponding protocols (CPD or air drying, respectively).

### Application to sEVs isolated from human serum

After having investigated the effects of different parameters on SEM imaging of NSCLC sEVs, we applied our optimized protocol to analyze and profile sEVs isolated from a human serum. The SEM protocol consisted of i) use of pure, filtered and not reactive chemicals, ii) covalent sEV capture to the substrate, iii) fixation and CPD, iv) Au/Pd sputtering. Furthermore, to demonstrate the prospect of using SEM for EV size profiling and as a means to check the purity of the EV isolation techniques, we isolated the vesicles by using two different methods, namely SEC and TFF. Figures 3A-D and Figures 3E-H show the pictures and SEM images of sEVs obtained from SEC and TFF based isolation methods, respectively. It has previously been reported that SEC allows for isolation of sEVs with minor contamination of proteins/small particles as compared to TFF.^26^ The latter cannot totally exclude small particles, despite being able to enrich samples within a tailored size window (in our case particles in the size range 30-200 nm). This can also be observed in the images presented in Figures 3, S6 and S7. As shown by the pictures of the collected samples (Figures 3A and 3E), the sEVs isolated by SEC showed a clear transparent appearance (Figure 3A), while those isolated by TFF from the same serum sample assumed a more yellowish and opaque aspect (Figure 3B). This was likely due to the presence of serum protein leftovers in the TFF sample. The SEM results also showed a clear difference among the two isolation methods. As presented in the representative SEM images in Figures 3, the substrate with sEVs from TFF method (Figures 3F-H and S7) showed a larger population of small particles in the size range of 10-50 nm than that obtained with the sEV sample isolated by SEC (Figures 3B-D and S6), for the same scanned areas.

**Figure 3.**
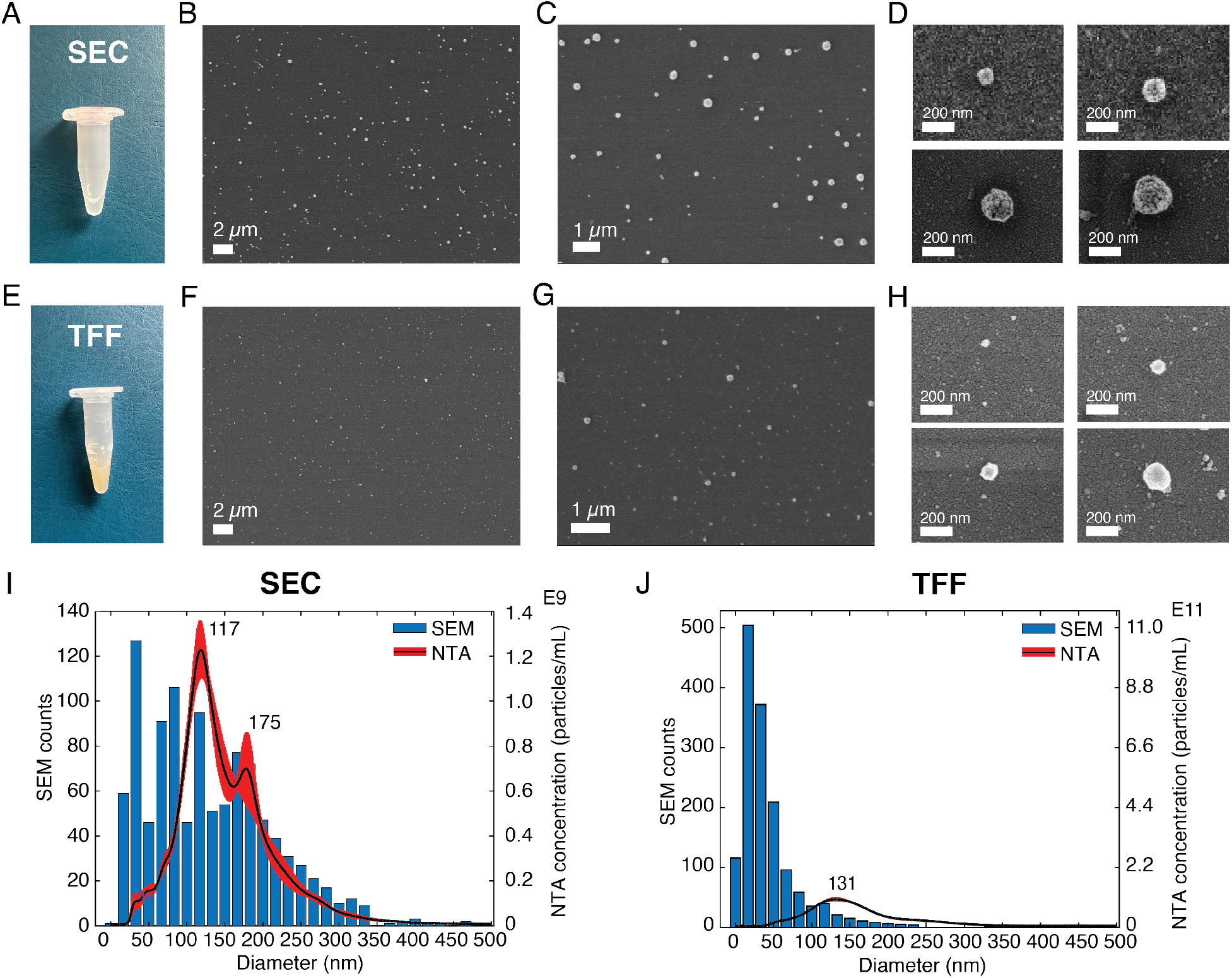
Validation of the SEM protocol on the sEVs isolated from a human serum. (A) Picture of the sEV sample isolated by using an IZON SEC kit. (B-D) Representative SEM images of the sEVs isolated by using SEC, showing vesicles of different diameters. AV=3kV for B-C, AV=1kV for D, for better vesicle surface resolution. (E) Picture of the sEV sample isolated by using the TFF technique. (F-H) Representative SEM images of the sEVs isolated by using TFF. AV=3kV for F-G, AV=1kV for H, for better vesicle surface resolution. (I) Comparison between the diameter distributions of the sEVs analyzed by SEM and those analyzed by NTA for the SEC method. (J) Comparison between the diameter distributions of the vesicles analyzed by SEM and those analyzed by NTA for the TFF method.

We further estimated the sEV size distributions by counting the vesicles in a large number of SEM images. Figures 3I and 3J show the histograms of representative size distribution obtained for the two sEV samples isolated by SEC and TFF, respectively. It can be seen that these size distributions also show the same qualitative behavior for the two different isolation methods. Unlike SEC, where the particle counts in the entire range of 10-200 nm were comparable, the TFF sample showed significantly larger particle counts below the size of ~70 nm. Furthermore, to validate our results we compared the particle distributions obtained by SEM with those obtained by NTA, the standard technique used for EV size analysis. The data clearly suggested a match between the distributions of the two methods for particles larger than 70 nm, while showing a difference for smaller particles. As presented and supported by other reports, the SEM accurately detected particles in the small size range (<50 nm) that could not be entirely detected by NTA or other scattering based methods.^5^ Here, we would also like to emphasize that the presence of a small shift between the two distributions, due to the characteristics of the techniques used, should be taken into account. While NTA measures the hydrodynamic radius of the particles and therefore it is likely to overestimate the size,^9^ the drying techniques needed for SEM may cause a shrinkage of the vesicles, leading to an underestimation of their diameters. In addition, the thin metal layer used for high resolution imaging may also cause an overall increase of the sizes detected by SEM.

## DISCUSSION

The importance of biophysical characterization and particularly size-based profiling of EVs have been addressed in a number of articles.^7,8,27^ Accurate size and abundance estimations of EVs remain a prerequisite for their clinical application, e.g. as a liquid biopsy source of biomarkers for cancers.^3,4,28^ A number of methods including NTA, TRPS and flow cytometry (FC) have been previously employed for this purpose. However, the results were found to be very different and technique dependent for the same EV sample.^5^ In biological samples, the variation in EV population may result in 25 fold differences in their sizes, 20 000 fold in volume and 10 000 000 fold in scattering intensity,^5^ meaning that a number of the established methods are accurate only for a fraction of the entire EV population range. For example, the size estimation by NTA and DLS relies on light scattering, which strongly depends on the particle size and material composition. In addition, interfacial properties, temperature and viscosity etc. also affects the results.^29^ Among the available techniques, electron microscopy still remains the most accurate approach which is also capable of covering the entire EV size range. In fact, a large number of reports on EVs have used TEM and SEM techniques but mainly as a means to visualize EVs rather than as a method for their systematic characterization and profiling. The technical bottleneck that has so far prevented these techniques to be a suitable EV size profiling method is their low throughput compared to other techniques such as NTA or FC. The results presented here aim to address this issue by methodically comparing different protocols and preparatory steps. As shown in Figure 1E, the throughput can be significantly improved by using a covalent capture of the vesicles to the substrates and by using a proper dehydration step (Figures 2B-C, 2E-F). These approaches also improve the EV distributions on the substrate, as seen in Figure 3B. Moreover, they allow for faster analysis within a short time frame (10-15 mins), as a sufficient number of images can be captured from different substrate areas for the EV size profiling.

The large measurable size range and nanometer scale accuracy of SEM also mean a better reliability of the measured size distributions. As presented in Figures 3I-J and as expected, compared to NTA the SEM based approach shows significantly larger EV counts in the size range below 70 nm, while closely following the NTA profile for larger diameters. This aspect has also been reported by other investigations^5,10^ and is mainly attributed to lower sensitivity and accuracy of NTA for smaller particles. In fact, depending on the preparation steps and methods, the EM-based approaches are also known to induce physical changes to the EV size and shapes.^25,30^ But such effects mainly introduce an overall shift of the distribution profile as all the vesicles irrespective of their sizes are affected in a similar manner. The major benefit of microscopy-based approaches is, however, their ability to accurately measure the population of the particles lying in the lower nanometer scale. As seen for the serum derived sEV sample (Figure 3), the size distribution measured by SEM extends to very small sizes of around 10 nm and a large number of particles were detected in that range. Although sEVs are not known to exist in such a small size, it is well known that a large variety of non-EV particles, e.g., protein aggregates, cell debris etc. may be co-isolated by the purification/isolation approach. These particles often produce significant challenges for downstream application/analysis.^31^ Their sizes are, however, far below the detection range of NTA and other scattering based approaches and therefore are not properly detected. The impurity may also include particles that have sizes similar to EVs. Previous studies have concluded that many EVs are spherical particles when in a physiological condition, while others have reported on different morphologies, such as oval or elongated vesicles.^32^ These morphological features of EVs should offer leverage to discriminate them from other non EV particles having random shapes, as shown in Figures 1G and 1I. Therefore, the SEM based approach can offer several benefits over the other approaches as the morphology and size can be monitored concomitantly. This advantage enables it to be a suitable method for both size-based EV profiling and optimization of the EV isolation/purification protocols.

Our investigation also clearly identifies different sources that can introduce artifacts which may be mistaken as EVs. These artifacts, within the scope of the present study, were mainly found to originate from the quality of reagents used and the process steps, as shown in Figures 1A, 1B and S2. Possibilities of such artifacts should be carefully evaluated in microscopic analysis of EVs. Furthermore, there has been several attempts to preserve the shape of EVs for microscopic studies in dry conditions, in particular by the use of glutaraldehyde and paraformaldehyde for fixation.^33^ Aldehyde reacts with free amino groups, stabilizing and cross linking the nucleic acid protein shell, and giving stability to the structure.^23,34^ However, as shown in Figures 2E-F and S5, CPD method seemed to work quite well for preserving the morphology of EVs and additional steps with GA/PFA did not show any significant improvement in our case.

## CONCLUSIONS

In conclusion, a SEM based approach for high resolution morphological analysis and size-based profiling of EVs is demonstrated. For this purpose, we developed a protocol for improved throughput, conformity, resolution and reproducibility of EV images by comparing various preparation and imaging steps. The optimized protocol was then used to profile the size distribution of sEVs derived from a human serum and also to compare the purity of two widely used EV isolation methods, i.e. size exclusion chromatography (SEC) and tangential flow filtration (TFF). The results revealed the presence of a higher number of small particles in the TFF sEV sample as compared to the SEC one. The size distribution profiles of these samples obtained by the proposed SEM based approach and the standard NTA based method were then compared for a qualitative assessment. The data revealed that while the distribution obtained by these two approaches closely followed each other for larger EVs (> 70 nm), they significantly deviated for smaller sizes. The SEM based approach clearly identified a larger proportion of EV like particles in smaller size range (< 70 nm). In addition, we also identified the presence of particles with random shapes within the reported EV size range from the analyzed EV samples, indicating the importance of morphological analysis for accurate EV identification and size profiling. In the future, we foresee that with some optimization, we can calibrate our SEM method to also obtain particle count estimation from an unknown sample.

## ASSOCIATED CONTENT

The following files are available free of charge. Supporting information containing the data on the characterization of the sEVs from NSCLC cells and human serum, as well as additional images of our different protocol combinations (PDF).

## AUTHOR INFORMATION

### Author Contributions

All authors have given approval to the final version of the manuscript.

SC conceptualized the study, developed the protocols, functionalized the substrates, performed the pre-imaging steps, acquired the SEM images, analyzed data, wrote original manuscript.

PH isolated the NSCLC H1975 sEVs and the sEVs from human serum using SEC, characterized all the sEV samples by NTA and WB.

KV edited and revised the manuscript, supervised the EV isolation and characterization by SEC and NTA.

AK isolated the sEVs from human serum using TFF, edited and revised the manuscript.

KF edited and revised the manuscript, acquired funding.

RL acquired funding.

JL edited and revised the manuscript, acquired funding.

AD conceptualized the study, wrote original manuscript, edited and revised manuscript and supervised the study.

### Conflicts of interest

There are no conflicts of interest to declare.

### Funding Sources

The supporting grants for this study were from the Erling Persson Family Foundation (to AD, JL, RL, KV and KF), Vatenskapsrådet (grant no. 2016-05051), the Stockholm Cancer Society ((#171123 (R.L) and #191293 (K.V)), the Swedish Cancer Society (CAN 2015/401; CAN 2018/597 to R.L), the Stockholm County Council (#20180404 to R.L) and Karolinska FOUU funding (#75032, #52614 (R.L. and K.V)).

## ACKNOWLEDGMENT

We would like to acknowledge Federico Pevere from Uppsala University (UU) for the help in sample sputtering and SEM imaging, and Victoria Sternhagen from UU for the suggestions and help in high resolution SEM imaging. MSc. Vasiliki Arapi from Karolinska Institutet is also acknowledged for her excellent help with western blot analyses.

## ABBREVIATIONS

EVs: extracellular vesicles
sEVs: small EVs
SEM: scanning electron microscopy
AFM: atomic force microscopy
NTA: nanoparticle tracking analyzer
FC: flow cytometry
SEC: size exclusion chromatography
TFF: tangential flow filtration.

## Notes

### Competing Interest Statement

The authors have declared no competing interest.

## REFERENCES

1 S. El Andaloussi, I. Mäger, X. O. Breakefield and M. J. A. Wood, Nat. Rev. Drug Discov., 2013, 12, 347–357.

2 D. E. Murphy, O. G. De Jong, M. Brouwer, M. J. Wood, G. Lavieu, R. M. Schiffelers and P. Vader, Exp. Mol. Med., 2019, 51, 1–12.

3 T. L. Whiteside, Expert Rev. Mol. Diagn., 2015, 15, 1293–1310.

4 R. Kalluri, J. Clin. Invest., 2016, 126, 1208–1215.

5 E. Van der Pol, F. A. Coumans, A. Grootemaat, C. Gardiner, I. Sargent, P. Harrison, A. Sturk,T. Van Leeuwen and R. Nieuwland, J. Thromb. Haemost., 2014, 12, 1182–1192.

6 E. Van der Pol, A. G. Hoekstra, A. Sturk, C. Otto, T. Van Leeuwen and R. Nieuwland, J. Thromb. Haemost., 2010, 8, 2596–2607.

7 F. Momen-heravi, L. Balaj, S. Alian, J. Tigges, V. Toxavidis, M. Ericsson, R. J. Distel, A. R. Ivanov, J. Skog and W. P. Kuo, Front. Physiol., 2012, 3, 1–8.

8 F. A. W. Coumans, A. R. Brisson, E. I. Buzas, F. Dignat-george, E. E. E. Drees, S. El-andaloussi, C. Emanueli, A. Gasecka, A. Hendrix, A. F. Hill, R. Lacroix, Y. Lee, T. G. Van Leeuwen, N. Mackman, I. Mäger, J. P. Nolan, E. Van Der Pol, D. M. Pegtel, S. Sahoo, P. R. M. Siljander, G. Sturk, O. De Wever and R. Nieuwland, Circ. Res., 2017, 120, 1632–1648.

9 D. Bachurski, M. Schuldner, P. Nguyen, A. Malz and K. S. Reiners, J. Extracell. Vesicles, DOI:10.1080/20013078.2019.1596016.

10 Y. Tian, M. Gong, G. Su, S. Zhu, W. Zhang, S. Wang, Z. Li, C. Chen, L. Li, L. Wu and X. Yan, ACS Nano, 2018, 12, 671–680.

11 Q. Zhang, J. N. Higginbotham, D. K. Jeppesen, Y. Yang, W. Li, T. Mckinley, R. Graves-deal, J. Ping, C. M. Britain, K. A. Dorsett, C. L. Hartman, D. A. Ford, R. M. Allen, K. C. Vickers, Q. Liu, L. Jeffrey, S. L. Bellis and R. J. Coffey, Cell Rep., 2019, 27, 940–954.

12 J. C. Akers, V. Ramakrishnan, J. P. Nolan, E. Duggan, C. Fu, F. H. Hochberg, C. C. Chen and B. S. Carter, PLoS One, 2016, 11, 1–11.

13 C. Lässer, S. Chul and L. Jan, Mol. Aspects Med., 2018, 60, 1–14.

14 M. Yáñez-Mó, P. R.-M. Siljander, Z. Andreu, A. Bedina Zavec and F. et al Borràs, J. Extracell. Vesicles, 2015, 4, 27066.

15 L. G. Rikkert, R. Nieuwland, L. W. M. M. Terstappen and F. A. W. Coumans, J. Extracell. Vesicles, , DOI:10.1080/20013078.2018.1555419.

16 L. Dubois, L. Löf, A. Larsson, K. Hultenby, A. Waldenström, M. Kamali-, G. Ronquist and K. G. Ronquist, Cancer Res. Front., 2018, 4, 13–26.

17 V. Sokolova, A. Ludwig, S. Hornung, O. Rotan, P. A. Horn, M. Epple and B. Giebel, Colloids Surfaces B Biointerfaces, 2011, 87, 146–150.

18 Y. Yang, G. Shen, H. Wang, H. Li, T. Zhang, N. Tao, X. Ding and H. Yu, 2018, 115, 10275–10280.

19 S. D. Ibsen, J. Wright, J. M. Lewis, S. Kim, S. Ko, J. Ong, S. Manouchehri, A. Vyas, J. Akers, C. C. Chen, B. S. Carter, S. C. Esener and M. J. Heller, ACS Nano, 2017, 11, 6641–6651.

20 C. Liu, X. Zeng, Z. An, Y. Yang, M. Eisenbaum, X. Gu, J. M. Jornet, G. K. Dy, M. E. Reid, Q. Gan and Y. Wu, ACS Sensors, 2018, 3, 1471–1479.

21 S. Cavallaro, J. Horak, P. Hååg, D. Gupta, C. Stiller, S. S. Sahu, A. Görgens, H. K. Gatty, K. Viktorsson, S. El Andaloussi, R. Lewensohn, A. E. Karlstrom, J. Linnros and A. Dev, ACS Sensors, 2019, 4, 1399–1408.

22 K. R. Jakobsen, B. S. Paulsen, R. Bæk, K. Varming, B. S. Sorensen and M. M. Jørgensen, J. Extracell. Vesicles, 2015, 4, 1–10.

23 Y. Chao and T. Zhang, Appl. Microbiol. Biotechnol., 2011, 92, 381–392.

24 M. Milani, D. Drobne and F. Tatti, Mod. Res. Educ. Top. Microsc., 2007, 787–794.

25 S. T. Chuo, J. C. Chien and C. P. Lai, J. Biomed. Sci., 2018, 25, 1–10.

26 G. Corso, I. Mäger, Y. Lee, A. Görgens, J. Bulte, B. Giebel, M. J. A. Wood, J. Z. Nordin and S. E. L. Andaloussi, Sci. Rep., 2017, 1–10.

27 J. C. Akers, V. Ramakrishnan, J. P. Nolan, E. Duggan, C. Fu, F. H. Hochberg, C. C. Chen and B. S. Carter, 2016, 1–11.

28 Z. Varga, B. Fehér, D. Kitka, A. Wacha, A. Bóta, S. Berényi, V. Pipich and J. Fraikine, Colloids Surfaces B Biointerfaces, 2020, 192, 111053.

29 C. M. Hoo, N. Starostin, P. West and M. L. Mecartney, J. Nanoparticle Res., 2008, 10, 89–96.

30 O. . Jensen, J. U. Prause and H. Laursen, Albr. von Graefes Arch. fur Klin. und Exp. Ophthalmol., 1981, 215, 233–242.

31 K. Sidhom, P. O. Obi and A. Saleem, Int. J. Mol. Sci., 2020, 21, 1–19.

32 H. Choi and J. Y. Mun, Appl. Microsc., 2017, 47, 171–175.

33 D. B. Nguyen, T. B. Thuy Ly, M. C. Wesseling, M. . Hittinger, A. . Torge, A. Devitt, Y. . Perrie and I. Bernhardt, Cell. Physiol. Biochem., 2016, 38, 1085–1099.

34 P. Parisse, I. Rago, L. U. Severino, F. Perissinotto, E. Ambrosetti, P. Paoletti, M. Ricci, A. Beltrami, D. Cesselli and L. Casalis, Eur. Biophys. J., 2017, 46, 813–820.

