## Supporting information for "High throughput imaging of nanoscale extracellular vesicles by scanning electron microscopy for accurate size-based profiling and morphological analysis"

**Figure S1**.

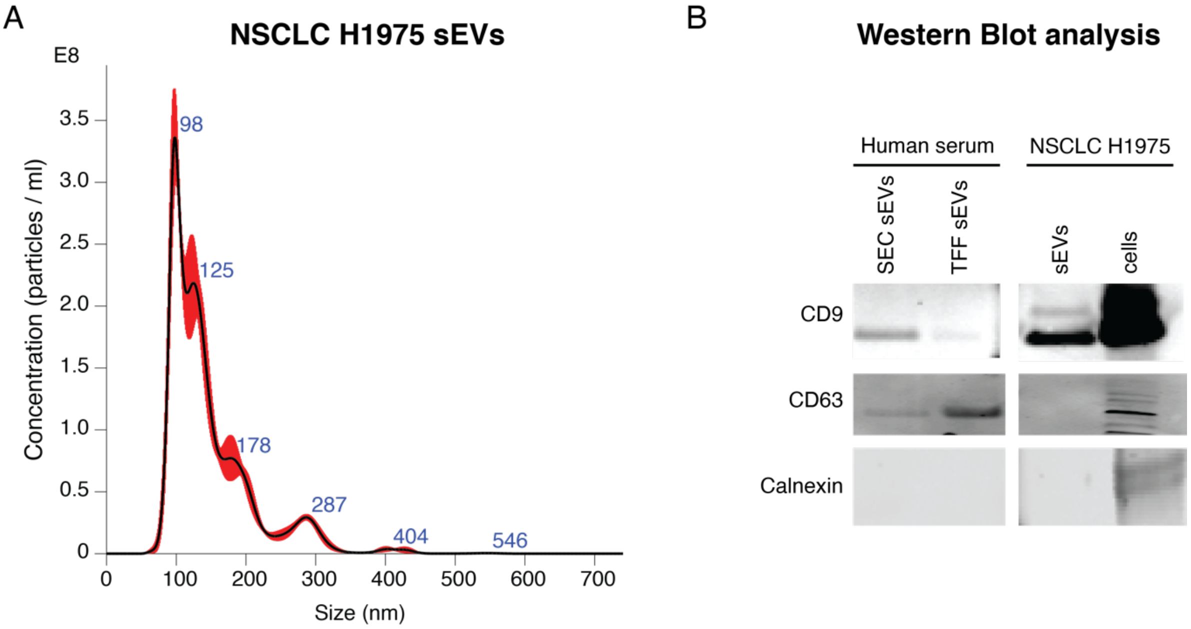


**Figure S1.** (A) NTA size distribution profile for the NSCLC H1975 sEVs. (B) Western Blot (WB) analysis of the NSCLC sEVs and the sEVs isolated from human serum by both SEC and TFF methods. As visible, the sEV samples showed presence of CD9 and CD63 tetraspanins and absence of calnexin, representative of cellular contamination. This was instead detected on NSCLC cells.

**Figure S2.**

A. No filtered chemicals and standard GA solution

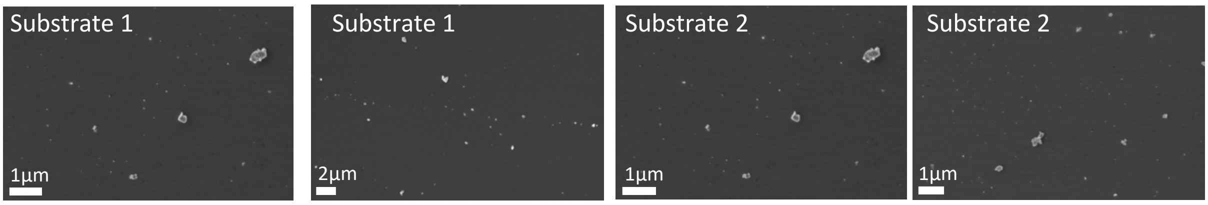

B. No deactivated GA

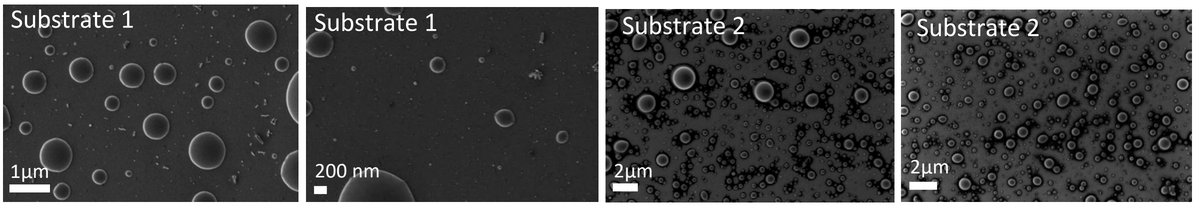


C. Filtered chemicals, Grade I GA solution and deactivated GA

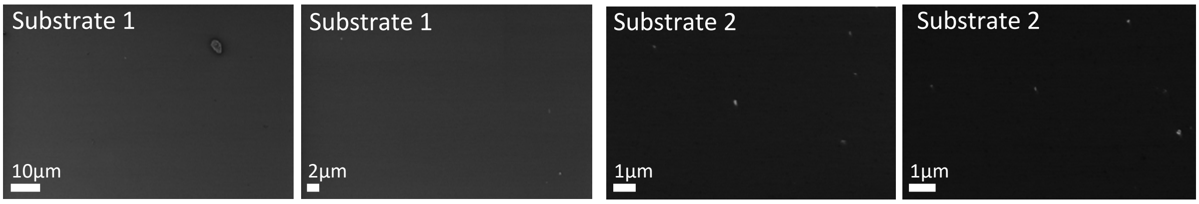


**Figure S2**. Representative control images of additional control substrates analyzed. (A) Substrates functionalized using as purchased chemicals (no filtered) and low-grade GA solution. AV=2kV. (B) Substrates functionalized following the control protocol up to the GA step. In this case, GA was not deactivated using Tris-ETHA and casein blocking agents. AV=2kV. (C) Good control substrates functionalized using filtered chemicals and Grade I and deactivated GA. AV=2kV. All substrates were dried using CPD.

**Figure S3**.

A. Non-covalent EV capture

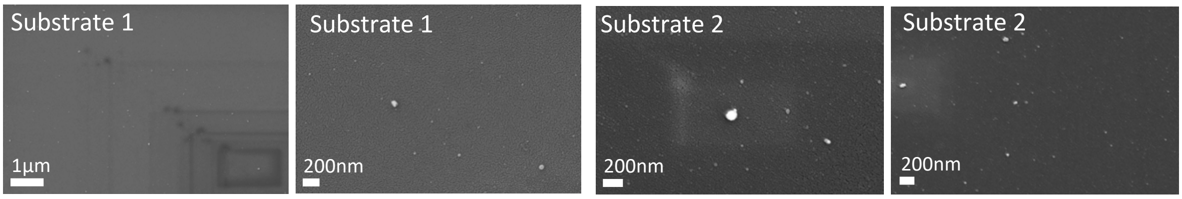


B. Covalent EV capture

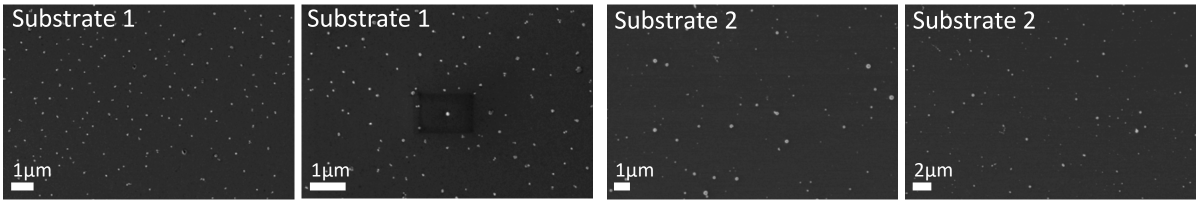


**Figure S3.** Representative images comparing substrates functionalized following the non-covalent vs covalent strategies. (A) Non-covalent EV capture. As shown, not many vesicles remained immobilized on the substrates. (B) Covalent EV capture. As shown, more vesicles remained immobilized on the substrates as compared to the non-covalent strategy, because of the stronger covalent binding as compared to the antibody-protein interaction.

**Figure S4**.

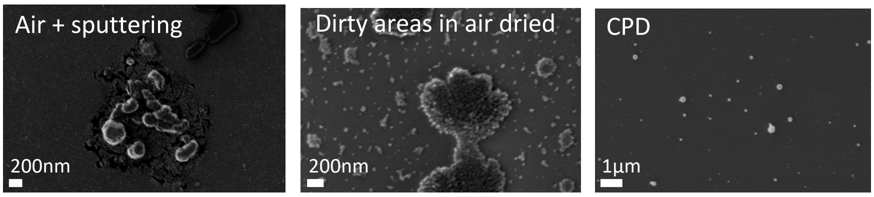


**Figure S4**. Representative images of additional substrates used to compare air drying with CPD. As shown, in the case of air drying, the wafers showed some dirty areas possibly arising from the drying process (middle image).

**Figure S5**.

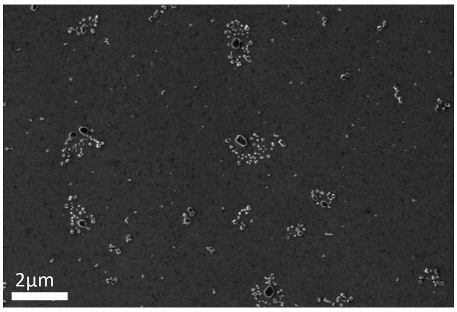


**Figure S5.** Representative image of a substrate with sEVs which was dried in air. In this case, the EVs were not fixed in the GA/PFA solution. As shown, the vesicles showed a similar shape and distribution as the substrates that were air dried with fixed vesicles (Figure 2).

**Figure S6**.

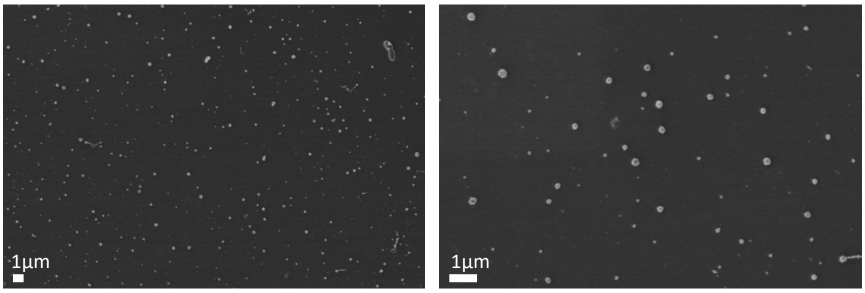

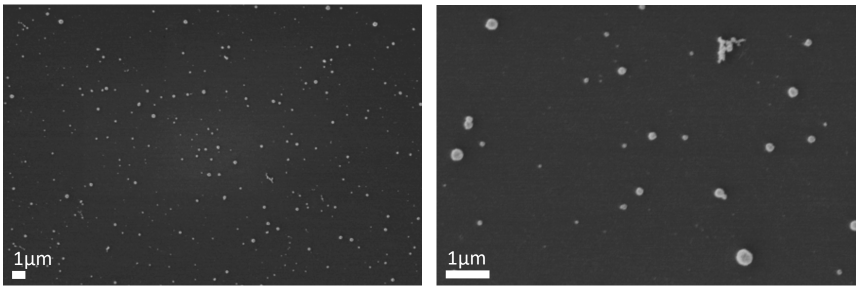


**Figure S6**. Additional representative images of the human serum sEVs isolated using the SEC method.

**Figure S7**.

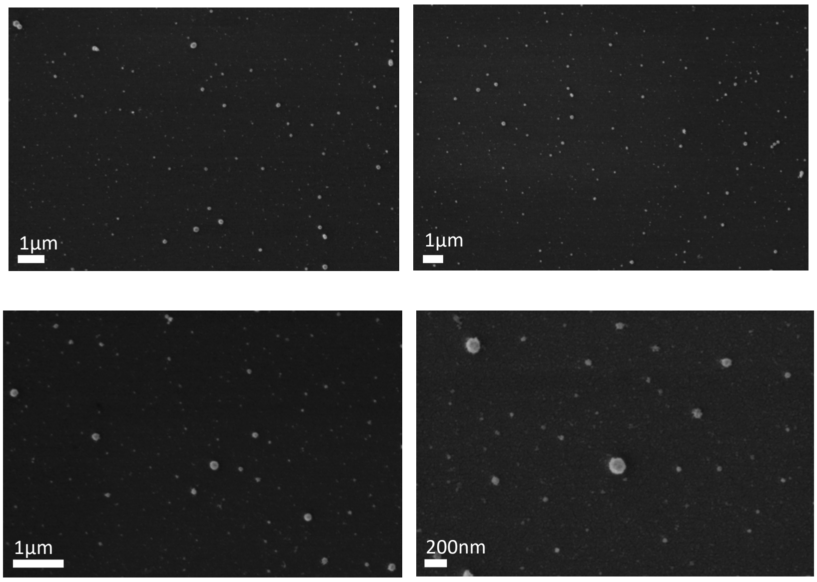


**Figure S7**. Additional representative images of the human serum sEVs isolated using the TFF method.
